## Supplementary Figures for "Multimodal determinants of phase-locked dynamics across deep-superficial hippocampal sublayers during theta oscillations"

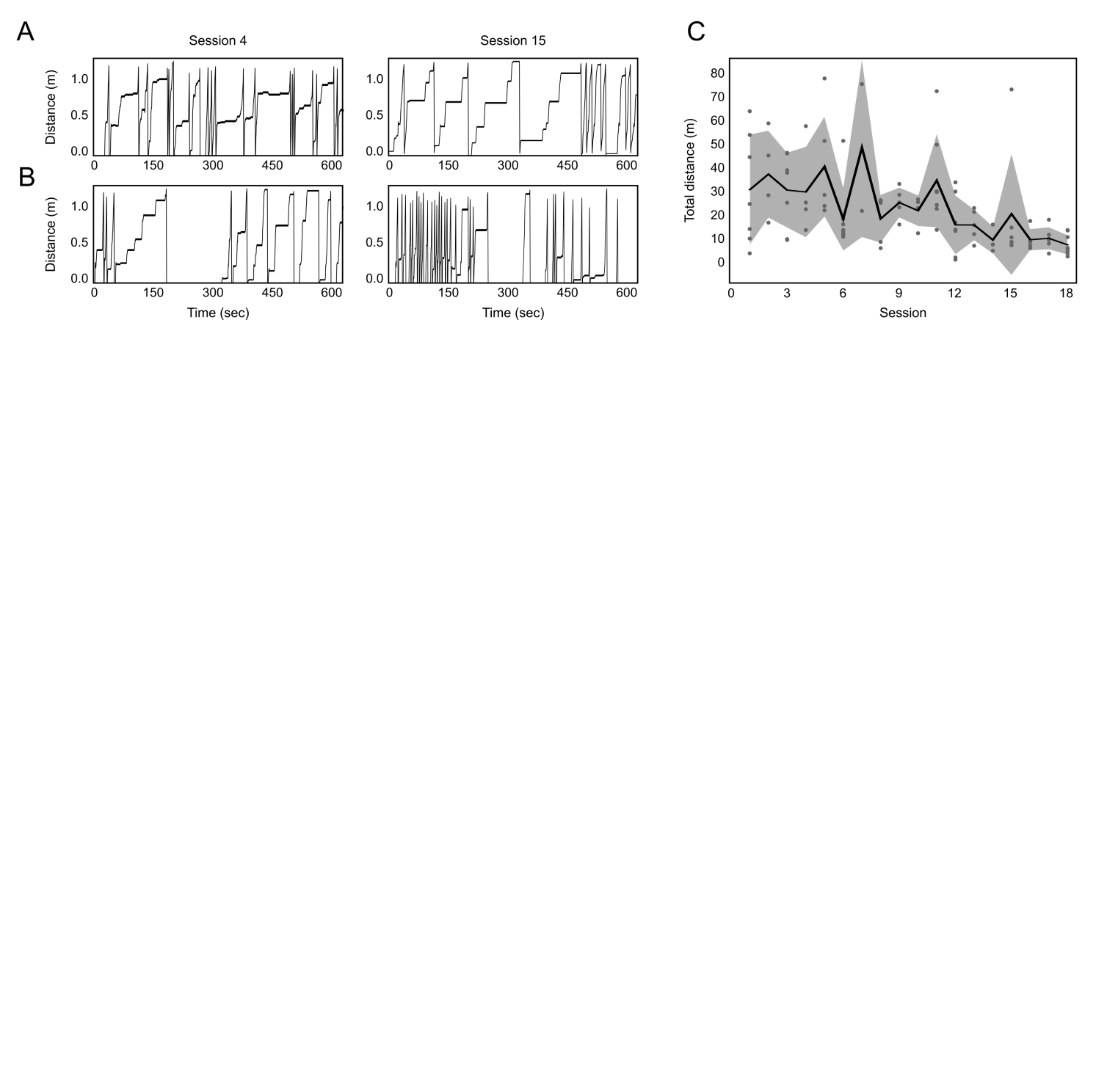

**Figure S1. Behavior of head-fixed mice. A**, Data from two sessions in one mouse showing habituation to the head-fixed setup. Animals run a minimum of 2 sessions of 10 min each per day over 2-3 weeks to get habituated to the setup before recordings start. The animal run freely without any particular stereotyped behavior, alternating between periods of running and immobility. During immobility periods, animals are attentive to the environment, they move their forelimbs for grooming or whisk. **B**, Example of sessions 4 and 15 from another mouse. **C**, Mean ± SD total distance traveled in 10 min sessions. Note animals habituate along sessions. Data from 18 sessions from 7 mice.

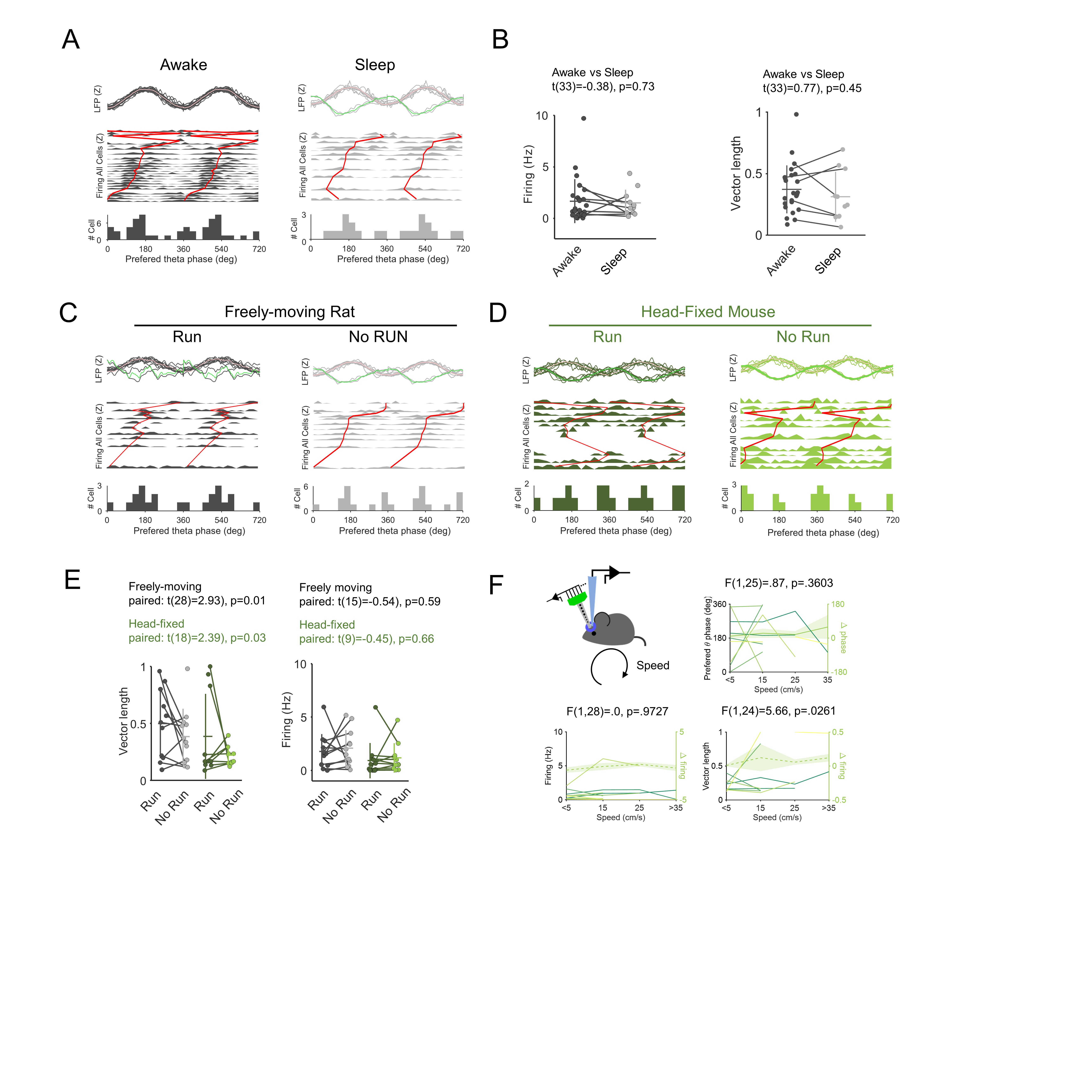

**Figure S2. Factors influencing theta phase-locked firing. A**, Data from freely moving rats were separated in theta episodes recorded during awake (n=25 cells) and REM sleep (n=11 cells). **B**, No state-dependent effect in data from freely moving rats (unpaired t-test). Data from cells recorded in both conditions are linked by lines (n=8 cells). **C**, Episodes of theta oscillations from freely moving rats were separated in periods of running (> 5 cm/sec; n=11 cells) versus periods of immobility (attentional or head-nodding theta; n=12 cells). **D**, Episodes of theta oscillations in head-fixed mice were separated in periods of wheel running (> 5 cm/sec; n=10 cells) versus theta periods during wheel immobility (attentional theta, whisking, forelimb movement; n=10 cells). **E**, Statistical effects of running versus no-running periods in both preparations. Data from cells recorded in both conditions are linked by lines (n=11 cells from rats, n=9 cells from mice). **F**, The effect of speed was examined in the head-fixed preparation (n=9 cells). We found some statistical effects for the mean vector length.

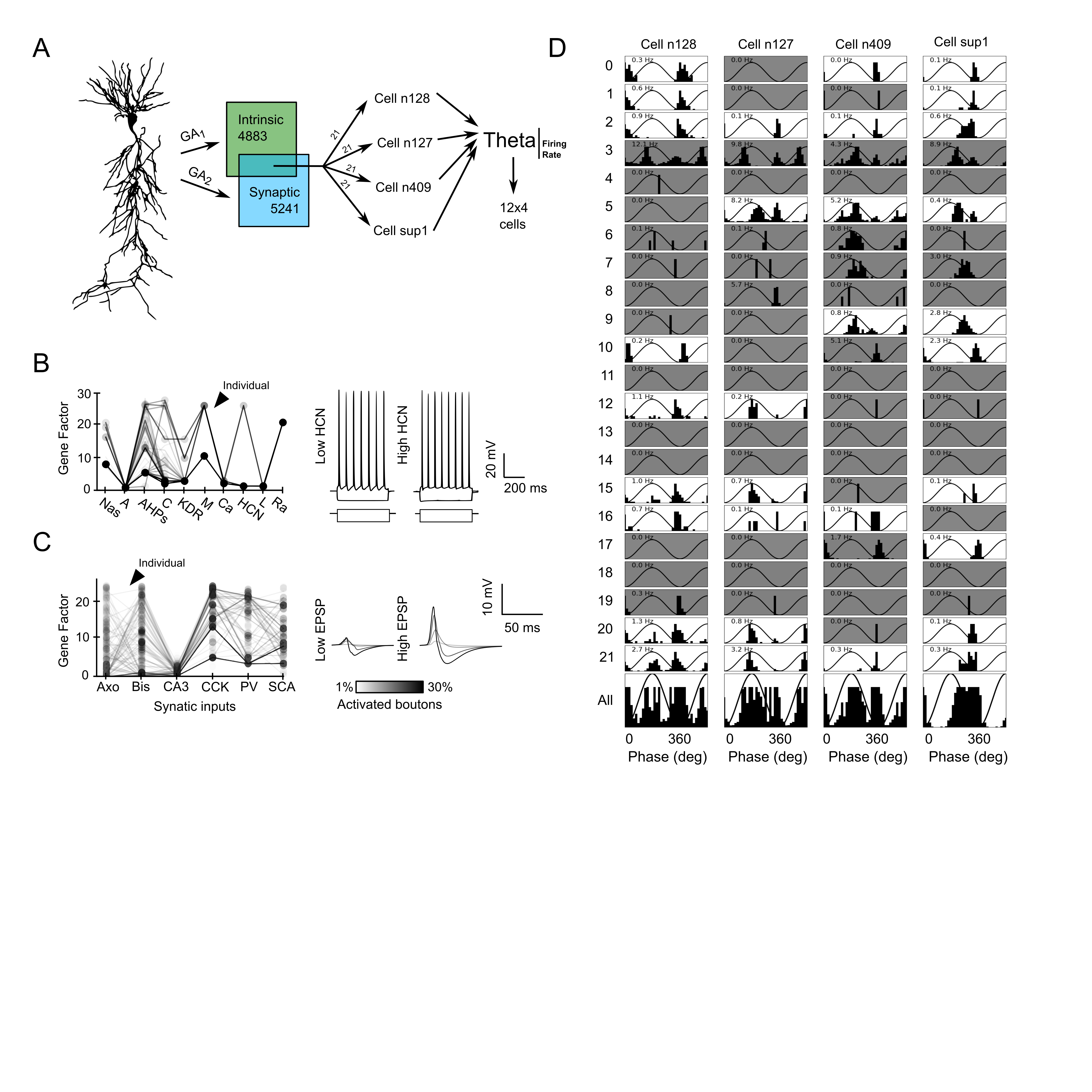

**Figure S3. Genetic algorithms and parameter constraints.** **A**, For a given morphology (cell n128) we used genetic algorithms (GA) to constrain intrinsic (ionic channel maximal conductances) and synaptic properties (maximal synaptic conductances) in a biologically realistic model of CA1 cells (Bezaire et al., 2016). Each GA provides a set of individuals (combination of genes) fitting experimental values for intrinsic and synaptic properties. We then chose 21 individuals fitted to both intrinsic and synaptic traits in four different morphologically-reconstructed CA1 pyramidal cells. These cells were submitted to a biologically realistic collection of glutamatergic (CA3, CA2, EC3, EC2) and GABAergic (Axo, Bis, CCK, Ivy, NGF, OLM, PV, SCA) theta-modulated inputs (450 cycles) impinging along different somatodendritic compartments. Theta modulation firing from each individual was evaluated and constrained by realistic rate values (0.1-5 Hz) yielding to 48 synthetic cells (12x4 morphologies). **B,** Solid lines link different combinations of genes giving experimentally valid input-output responses shown in Fig.2B. Arrowhead point to one individual. Examples of current pulse responses by two different individuals are shown at right. **C**, Same as in B for fitting synaptic responses to Schaffer collaterals stimulation with two individual examples. **D**, Theta phase locked firing of the different synthetic cells. Gray shadowed cells did not meet realistic rate values.

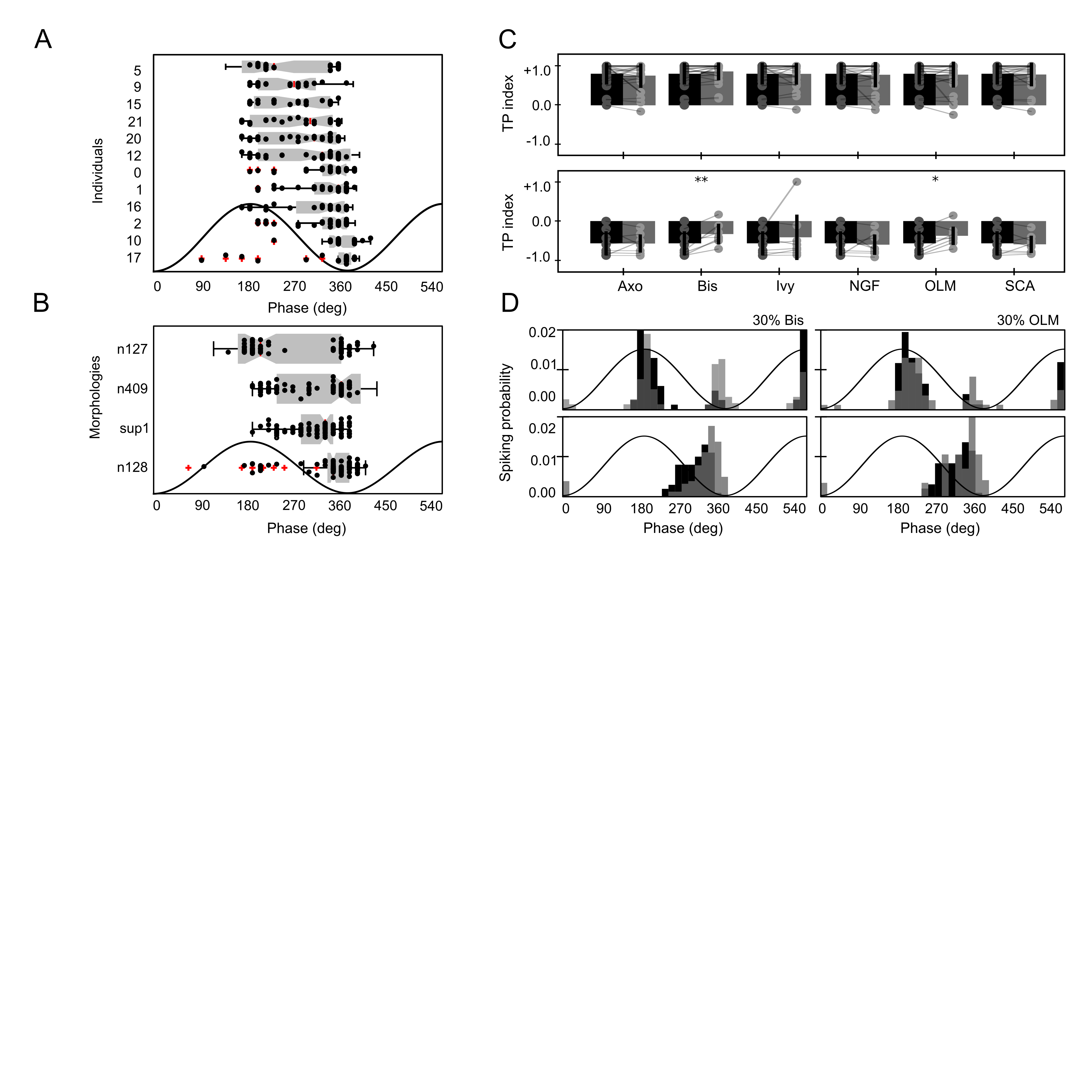

**Figure S4. Effect of GABAergic connectivity on theta phase preference. A**, Theta phase distribution across individuals with different PV/CCK connectivity (n=48 synthetic cells). Individuals are ranked according to the order in Fig.2H. **B**, Effect of morphology in the distribution of preferred theta phases caused by different PV/CCK connectivity. Cell morphologies are ranked according to the order in Fig.2G. **C**, Effect of 70% reduction of GABAergic inputs from different interneuronal types. Statistical differences were confirmed only for bistratified and OLM synthetic cells. *, p<0.05; **, p<0.01. **D**, Representative examples of the effect of 30% reduction of GABAergic inputs from bistratified and OLM cells in synthetic cells tuned to the theta peak (upper row) and trough (lower row).

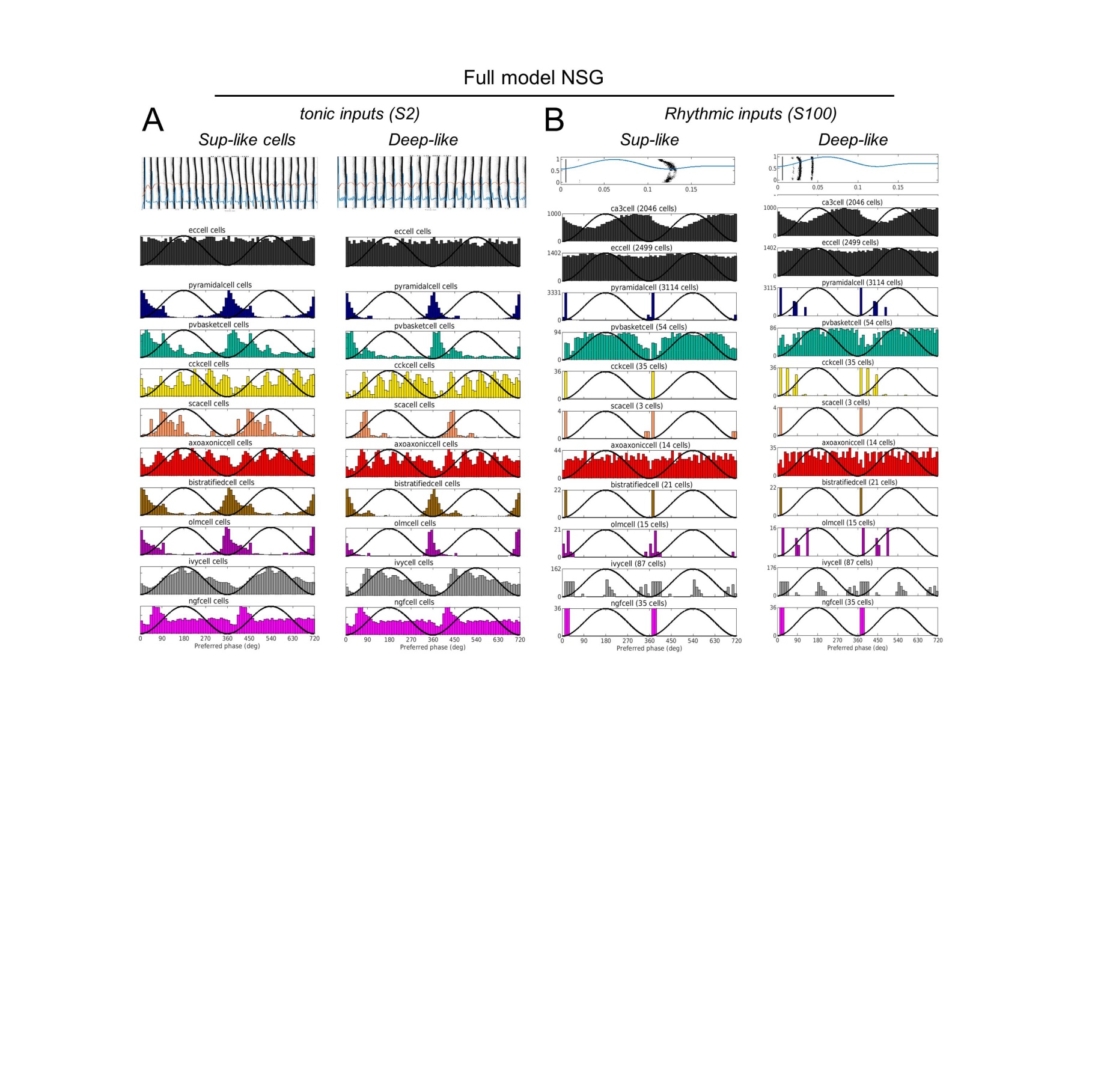

**Figure S5. Comparison with the full model.** **A**, Simulation results of superficial- and deep-like parameters in the full model (scale 2) run at the Neuroscience Gateway (NSG) supercomputers. Note that this model generates theta oscillations autonomously in response to tonic inputs. Using deep- and superficial-like PV/CCK connectivity yielded theta oscillations at similar frequencies: 12 Hz for superficial-like cells and 11 Hz for deep-like cells. Since the LFP signal in the full model is calculated from pyramidal cell activity, the effect of phase-locking preference cannot be tested. **B**, Simulation results of superficial- and deep-like parameters in the full model (scale 100) for rhythmic inputs. In this simulation glutamatergic CA3 and EC3 inputs were theta-modulated to provide with a general clock and LFP-like theta signal. Using superficial- and deep-like connectivity resulted in different phase-locking preference, similar to our simulation results. Note that phase-locked firing by PV and CCK basket cells differ from the original simulations.

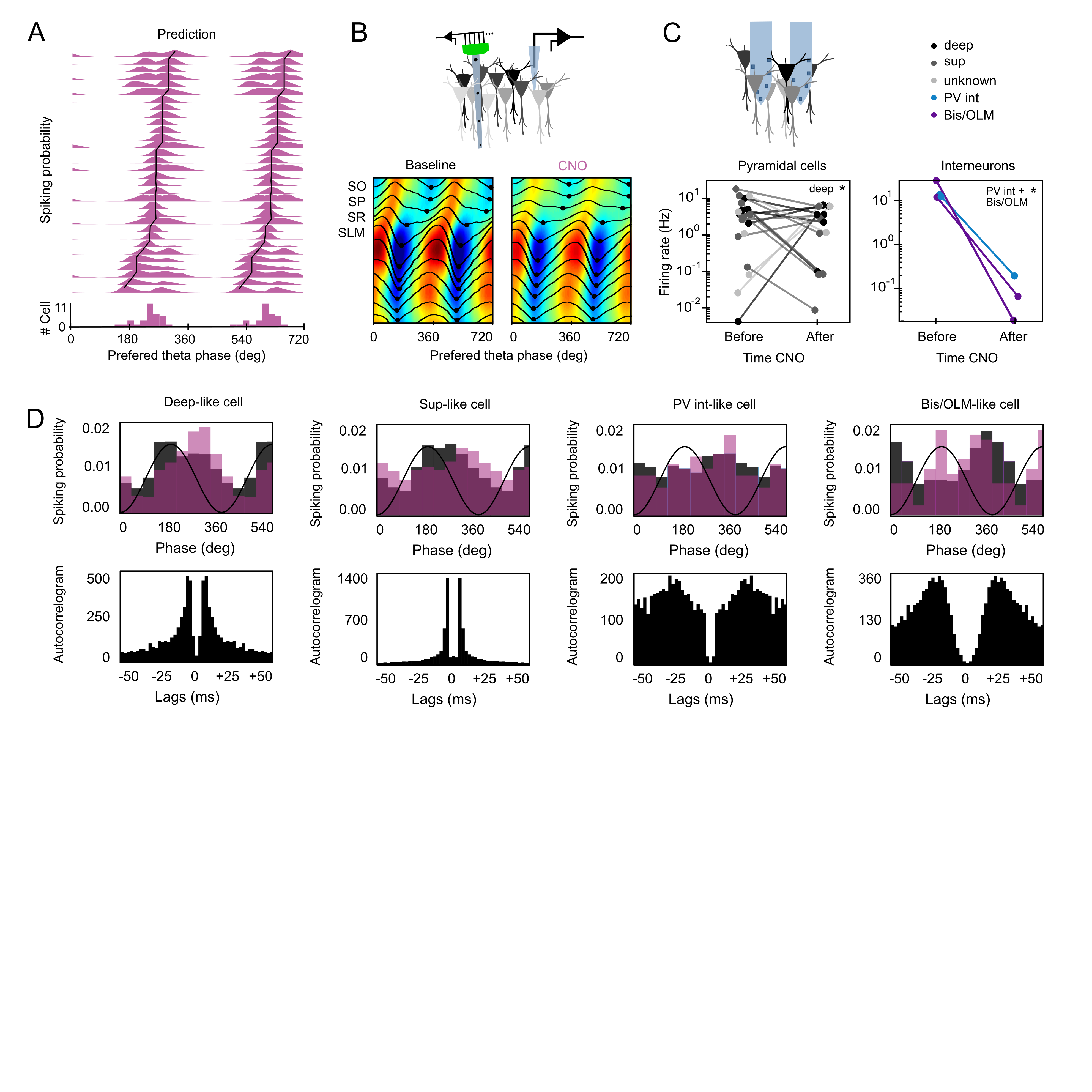

**Figure S6. Effects of chemogenetic silencing of PV cells. A,** Model prediction of the population level effect of blocking PV basket cell inputs in synthetic cells (n=32). **B**, Similar laminar profiles of LFP and CSD theta cycles with respect to SLM after i.p. CNO injection in PV-Cre mice injected with AAV8-DIO-hSyn-hM4D(Gi)-mCherry. **C**, Multisite silicon probes allowed to evaluate changes of firing rate before and after CNO within putatively identified cell types (deep and superficial pyramidal cells and PV-, bistratified, OLM-like interneurons). Note significantly reduction of PV- and Bis/OLM-like cells, consistent with anatomical confirmation of hM4D(Gi) expression. **D**, Theta-phase firing histograms and autocorrelation before and after CNO for different cell types.

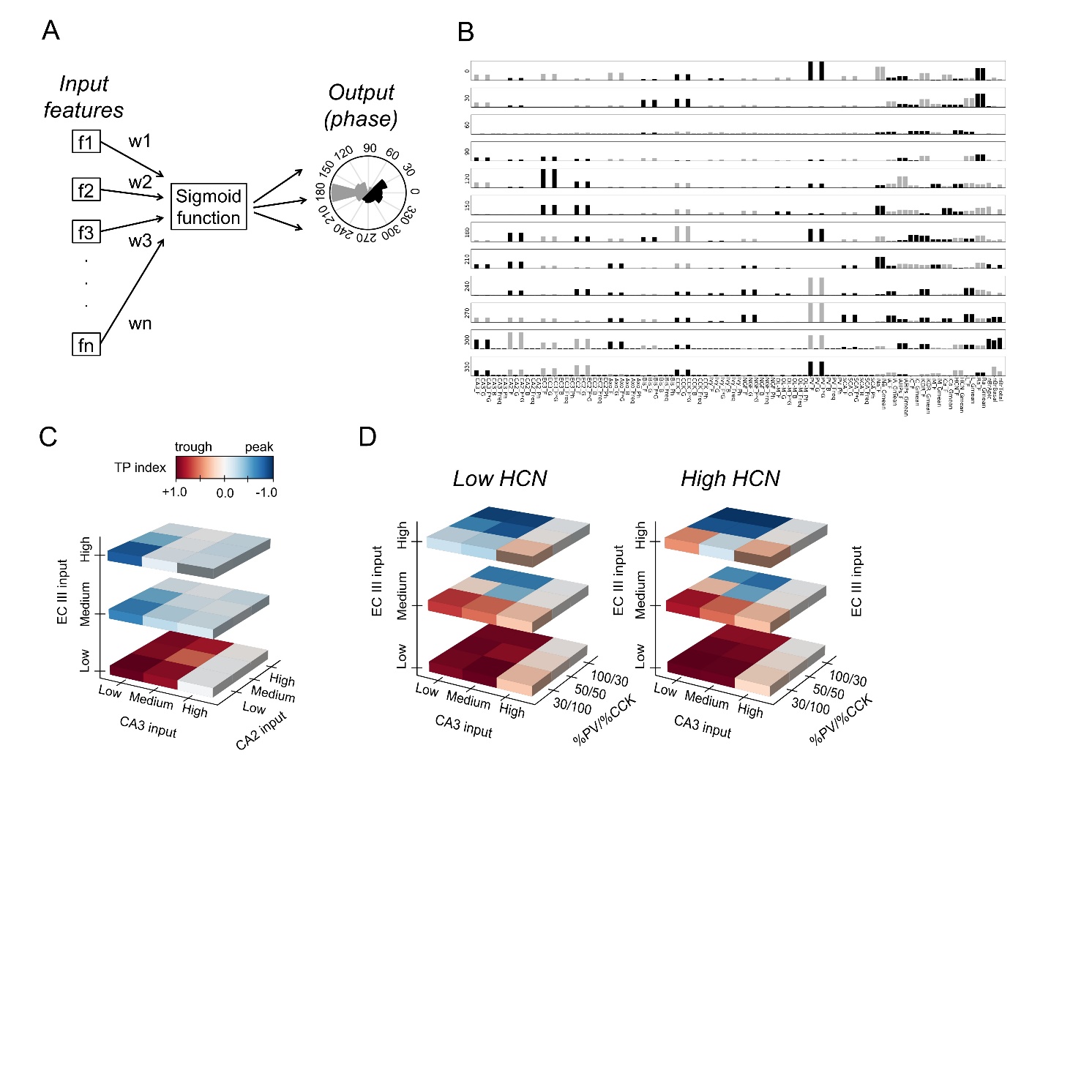

F**igure S7. Logistic regression model. A**, A multinomial logistic regression model was implemented to evaluate the relative contribution of different biophysical, intrinsic and connectivity factors (97 factors) in determining phase timing using 731 heterogeneous synthetic cells. **B**, Subset of all significant contributing factors out of the 97 features examined. Black represents the effect of upregulation, gray represents the effect of downregulation. **C**, Effect of the interaction between CA3, ECIII and CA2 input pathways in phase shifting a single individual cell. **D**, Effect of low and high levels of HCN in phase timing individual cells to CA3 and ECIII input pathways as a function of the PV/CCK axis.

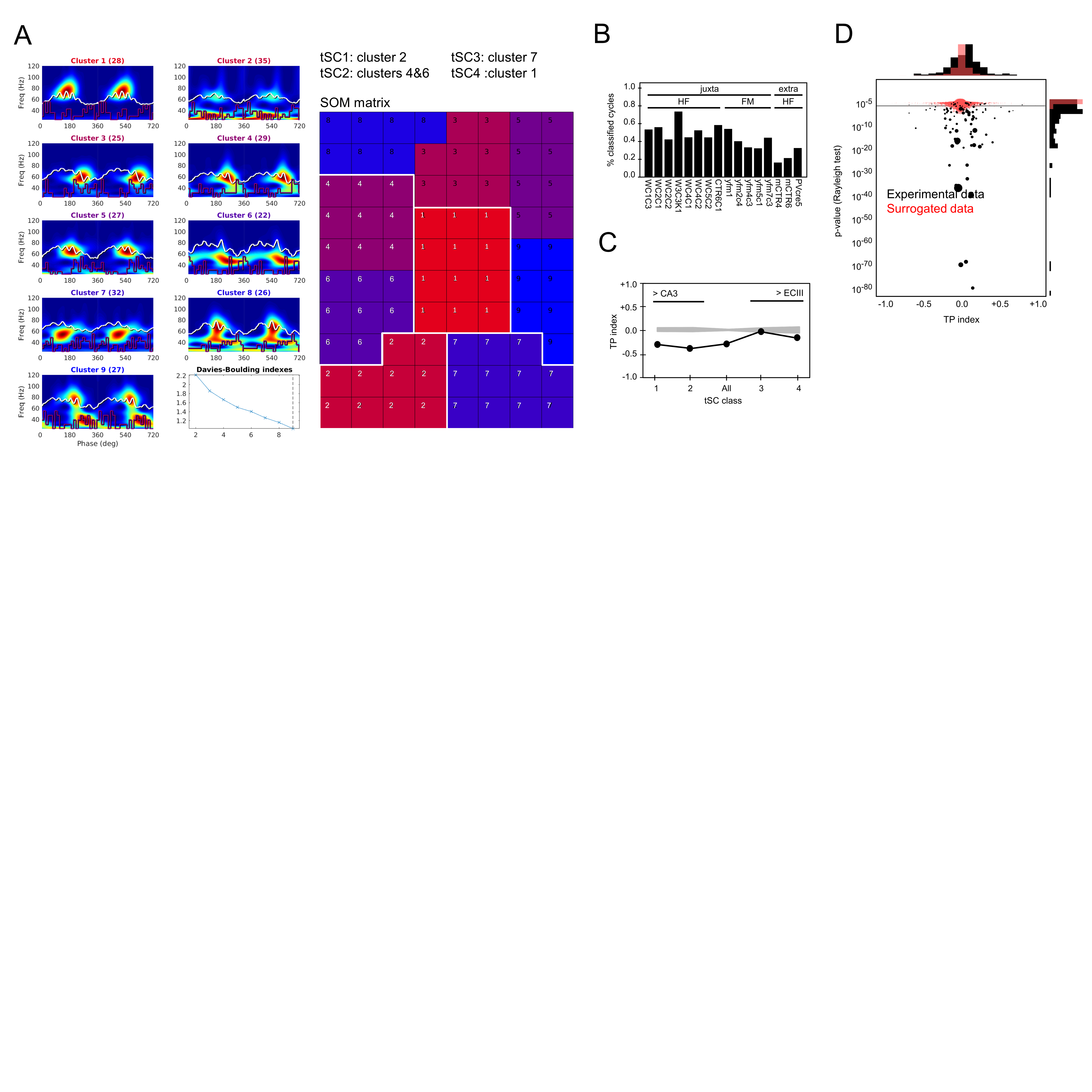

**Figure S8. Effect of input pathways on theta phase-locked firing. A**, Clusters of individual theta cycles classified automatically by SOM are shown together with their mean time-frequency spectra. Number in parentheses refer to the number of individual cycles at each cluster. The SOM matrix at left shows the topological relationship between clusters. Theta nested spectral components are blind to SOM. However, since the algorithm clustered cycles based on their waveforms, the different spectral classes (tSC1, 2, 3 and 4) are successfully identified. Note that clusters 4 and 6 were assigned to class tSC2 given their topological proximity and comparable spectral components. **B**, Percentage of classified theta cycles from the different experiments. HF, head-fixed; FM, freely moving. **C,** Comparison between TP index values estimated for each tSC class and those obtained from 1000 surrogates (gray) for the cell shown in Fig.4. **D,** Results from the surrogate tests (red; 1000 realizations) on the experimental data (black).

**Table S1 Conductance model parameters taken from literature.**

| **Channel** | **Location** | **Distance to soma (um)** | **Conductance (S/cm^2^)** | **Reference** |
| --- | --- | --- | --- | --- |
| iA | Soma | all | 0.0025 | Cutsuridis (2015) NLM. ModelDB #181967 |
| iA | Basal | all | 0.0600 |  |
| iA | Apic | 50 | 0.0600 |  |
| iA | Apic | 200 | 0.0600 |  |
| iA | Apic | 350 | 0.0600 |  |
| iAHPs | Soma | all | 0.0005 | Cutsuridis (2015) NLM. ModelDB #181967 |
| iAHPs | Basal | all | 0.0005 |  |
| iAHPs | Apic | 50 | 0.0005 |  |
| iAHPs | Apic | 200 | 0.0005 |  |
| iAHPs | Apic | 350 | 0.0005 |  |
| iC | Soma | all | 0.09075 | Cutsuridis (2015) NLM. ModelDB #181967 |
| iC | Basal | all | 0.03300 |  |
| iC | Apic | 50 | 0.03300 |  |
| iC | Apic | 200 | 0.03300 |  |
| iC | Apic | 350 | 0.00410 |  |
| iCaL | Soma | all | 0.000700000 | Mueller et al. (2015) Neuron. ModelDB #206244 |
| iCaL | Basal | all | 0.000031635 |  |
| iCaL | Apic | 50 | 0.000031635 |  |
| iCaL | Apic | 200 | 0.000031635 |  |
| iCaL | Apic | 350 | 0.000031635 |  |
| iCaT | Soma | all | 0.00005 | Mueller et al. (2015) Neuron. ModelDB #206244 |
| iCaT | Basal | all | 0.00001 |  |
| iCaT | Apic | 50 | 0.00001 |  |
| iCaT | Apic | 200 | 0.00001 |  |
| iCaT | Apic | 350 | 0.00001 |  |
| iKDR | Soma | all | 0.001400 | Cutsuridis (2015) NLM. ModelDB #181967 |
| iKDR | Axon | all | 0.020000 |  |
| iKDR | Basal | all | 0.000868 |  |
| iKDR | Apic | 50 | 0.000868 |  |
| iKDR | Apic | 200 | 0.000868 |  |
| iKDR | Apic | 350 | 0.000868 |  |
| iKDR | Apic | 800 | 0.000868 |  |
| iM | Soma | all | 0.06 | Cutsuridis (2015) NLM. ModelDB #181967 |
| iM | Axon | all | 0.03 |  |
| iM | Basal | all | 0.06 |  |
| iM | Apic | 50 | 0.06 |  |
| iM | Apic | 200 | 0.06 |  |
| iM | Apic | 350 | 0.06 |  |
| iNas | Soma | all | 0.07 | Cutsuridis (2015) NLM. ModelDB #181967 |
| iNas | Axon | all | 0.07 |  |
| iNas | Basal | all | 0.07 |  |
| iNas | Apic | 50 | 0.07 |  |
| iNas | Apic | 200 | 0.07 |  |
| iNas | Apic | 350 | 0.07 |  |
| iNas | Apic | 800 | 0.07 |  |
| iHCN | Apic | all |  | Sinha (2015) PNAS |

**Table S2 Synaptic model parameters taken from literature.** * Bezaire et al. (2016), Bezaire et al. (2013), Klausberger and Somogyi (2008), Mizuseki and Buzsaki (2013)

| **Pre-synaptic Cell** | **# boutons** | **Location** | | **Distance to soma (um)** | **Frequency (Hz)** | **E_rev_ (mV)** | **ττ_1_ (ms)** | **ττ_2_ (ms)** | **G_max_** | **% Basal firing** | **Phase (deg)** | **β_1_** | **β_2_** | **References*** |
| --- | --- | --- | --- | --- | --- | --- | --- | --- | --- | --- | --- | --- | --- | --- |
| CA3 | 6209 | Apic | SLM | 50 to 300 | 1.5 | 0 | 0.5 | 3 | 0.0002 | 0.5 | 276 | 5 | 3 |  |
| CA3 | 2661 | Apic | SR (thick) | 50 to 300 | 1 | 0 | 0.5 | 3 | 0.0002 | 0.5 | 276 | 5 | 3 |  |
| CA3 | 34 | Apic | SLM | 300 to 600 | 1 | 0 | 0.5 | 3 | 0.0002 | 0.5 | 276 | 5 | 3 |  |
| CA3 | 3000 | Axon | SP | -200 to 0 | 1 | 0 | 0.5 | 3 | 0.0002 | 0.5 | 276 | 5 | 3 |  |
| CA2 | 8000 | Basal | SO | -200 to 0 | 1 | 0 | 0.5 | 3 | 0.0002 | 0.25 | 330 | 5 | 5 |  |
| EC3 | 968 | Apic | SR (thick) | 300 to 600 | 1 | 0 | 0.5 | 3 | 0.0002 | 0.8 | 180 | 3 | 5 |  |
| EC3 | 774 | Apic | SLM | 300 to 600 | 1.5 | 0 | 0.5 | 3 | 0.0002 | 0.8 | 180 | 3 | 5 |  |
| EC2 | 700 | Apic | SR (thick) | 300 to 600 | 1 | 0 | 0.5 | 3 | 0.0002 | 0.8 | 330 | 5 | 5 |  |
| EC2 | 1042 | Apic | SLM | 300 to 600 | 1.5 | 0 | 0.5 | 3 | 0.0002 | 0.8 | 330 | 5 | 5 |  |
| Axo | 34 | Axon | SP | -200 to 0 | 20 | -70 | 0.28 | 8.4 | 0.00115 | 0 | 185 | 10 | 6 | Forro et al. (2013) |
| Bis | 53 | Basal | SO | -200 to -50 | 40 | -70 | 0.11 | 9.70 | 0.00051 | 0 | 1 | 5 | 5 | Bulh et al. (1996) |
| Bis | 3 | Basal | SO | -50 to 0 | 40 | -70 | 0.11 | 9.70 | 0.00051 | 0 | 1 | 5 | 5 |  |
| Bis | 4 | Apic | SR (thin) | 0 to 150 | 40 | -70 | 0.11 | 9.70 | 0.00051 | 0 | 1 | 5 | 5 |  |
| Bis | 44 | Apic | SR (thin) | 0 to 300 | 40 | -70 | 0.11 | 9.70 | 0.00051 | 0 | 1 | 5 | 5 |  |
| CCK | 37 | Basal | SO | -50 to 0 | 60 | -70 | 0.2 | 4.2 | 0.00052 | 0.25 | 174 | 3 | 5 | Mátyás et al. (2004) |
| CCK | 37 | Apic | SR (thick) | 0 to 150 | 60 | -70 | 0.2 | 4.2 | 0.00052 | 0.25 | 174 | 3 | 5 |  |
| CCK | 30 | Soma | SP | -10 to 10 | 60 | -70 | 0.2 | 4.2 | 0.00052 | 0.25 | 174 | 3 | 5 |  |
| CCK | 1 | Apic | SR (thick) | 300 to 600 | 60 | -70 | 0.2 | 4.2 | 0.00052 | 0.25 | 174 | 3 | 5 |  |
| Ivy | 169 | Basal | SO | -200 to -50 | 4 | -70 | 1.1 | 11 | 0.000041 | 0.5 | 31 | 3 | 5 | Fuentealba et al. (2008) |
| Ivy | 4 | Basal | SO | -50 to 0 | 4 | -70 | 1.1 | 11 | 0.000041 | 0.5 | 31 | 3 | 5 |  |
| Ivy | 4 | Apic | SR (thin) | 0 to 150 | 4 | -70 | 1.1 | 11 | 0.000041 | 0.5 | 31 | 3 | 5 |  |
| Ivy | 211 | Apic | SR (thin) | 0 to 300 | 4 | -70 | 1.1 | 11 | 0.000041 | 0.5 | 31 | 3 | 5 |  |
| Ivy | 34 | Apic | SLM | 300 to 600 | 4 | -70 | 1.1 | 11 | 0.000041 | 0.5 | 31 | 3 | 5 |  |
| NGF | 24 | Apic | SR (thin) | 0 to 300 | 4 | -70 | 9 | 39 | 0.000065 | 0 | 196 | 4 | 8 | Suzuki et al. (2014) |
| NGF | 116 | Apic | SLM | 300 to 600 | 4 | -70 | 9 | 39 | 0.000065 | 0 | 196 | 4 | 8 |  |
| OLM | 5 | Basal | SO | -200 to -50 | 23 | -70 | 0.13 | 11 | 0.0003 | 0.1 | 19 | 6 | 6 | Scorza et al. (2011) |
| OLM | 72 | Apic | SLM | 300 to 600 | 23 | -70 | 0.13 | 11 | 0.0003 | 0.1 | 19 | 6 | 6 |  |
| PV | 61 | Basal | SO | -50 to 0 | 50 | -70 | 0.3 | 6.2 | 0.0002 | 0.3 | 271 | 15 | 10 | Lasztóczi and Klauberger (2014) |
| PV | 61 | Apic | SR (thick) | 0 to 150 | 50 | -70 | 0.3 | 6.2 | 0.0002 | 0.3 | 271 | 15 | 10 |  |
| PV | 61 | Soma | SP | -10 to 10 | 50 | -70 | 0.3 | 6.2 | 0.0002 | 0.3 | 271 | 15 | 10 |  |
| SCA | 1 | Basal | SO | -200 to -50 | 7 | -70 | 0.3 | 8 | 0.00037 | 0 | 205 | 5 | 5 |  |
| SCA | 12 | Apic | SR (thin) | 0 to 300 | 7 | -70 | 0.3 | 8 | 0.00037 | 0 | 205 | 5 | 5 |  |
| SCA | 1 | Apic | SR (thick) | 300 to 600 | 7 | -70 | 0.3 | 8 | 0.00037 | 0 | 205 | 5 | 5 |  |
